# Butterfly brains change in morphology and in gene splicing patterns after brief pheromone exposure

**DOI:** 10.1101/2024.10.02.615994

**Authors:** Emilie Dion, YiPeng Toh, Dantong Zhu, Antónia Monteiro

## Abstract

How insect brains differ between the sexes and respond to sex-specific pheromones is still not well understood. Here we briefly exposed female *Bicyclus anynana* butterflies to wild type (Wt) and modified male sex pheromone blends, previously shown to modify females’ sexual preferences, and examined how their brains were modified at the morphological and molecular levels, three days later. First, we 3D-reconstructed male and female brains of this species and documented sexual dimorphism in the size of seven of 67 glomeruli present in the olfactory lobe. Then we showed that several glomeruli changed in volume after blend exposures, implicating them in sex pheromone perception. Finally, we found that a few genes were differentially expressed but many more were differentially spliced between male and female naïve brains, and between naive and pheromone blend-exposed brains. These are primarily calcium-binding channels and RNA-binding genes, respectively. A learned preference for changed levels in a single pheromone component was linked to variants of proteins involved in synaptic transmission. Our work shows that naïve male and female brains differ primarily in gene splicing patterns and that a brief, 3-minute, exposure to pheromones produces slight changes in brain volume and large changes in the splicing of genes involved in neural development, that correlate with changes in sexual preferences in females.

**Significance statement:** How brains differ between the sexes and respond to sex-specific cues is a hot research topic. Here we investigate how the brains of female butterflies differ from those of males and respond to male sex pheromones. We find that the sexes differ in the volume of a sub-set of olfactory lobe glomeruli, and the volume of some glomeruli also changes after exposure to pheromone blends. In addition, male and female brains differ primarily in hundreds of splice variants, both before and after pheromone exposure. These findings suggest that different proteins (splice variants) characterize male and female brains and that a brief exposure to pheromones can lead to changes in brain structure and in further gene splicing linked to altered sexual preferences in female butterflies.

## Introduction

Odour learning in insects can impact sex pheromone preferences and the evolution of populations and species. For instance, individuals of several species can learn to prefer new odours via exposure to those blends (Adam et al., 2022; Fabian and Sachse, 2023). If these odours are novel sex pheromone blends, populations that evolve a preference for these blends can potentially undergo rapid reproductive isolation and speciation (Dion et al., 2019; Zhao and McBride, 2020; Anton and Rössler, 2021). It is, thus, interesting to investigate how pheromone odour learning might take place at the mechanistic level.

Odour learning is a plastic behavioural response that affects all levels of the olfactory circuitry. It can be measured and examined using different physiological and molecular tools. For instance, mechanisms of chemosensory learning have been explored with electrophysiology experiments at the periphery and in the central nervous system, via the measurement of neuronal activities (e.g. Anderson et al. (2007); Arenas et al. (2009); Guerrieri et al. (2012); López et al. (2017); Seeholzer et al. (2018)). In addition, these mechanisms have been explored at the molecular level in the insect peripheral sensory structures, such as antennae and legs, via monitoring of molecular changes after an odour exposure (e.g. Claudianos et al. (2014); Wan et al. (2015)). However, the molecular actors of olfactory learning in the insect’s central nervous system remain underexplored.

The antennal lobes (ALs) and the mushroom bodies (MBs) are the main brain structures involved in olfactory learning (Stopfer, 2014; Amin and Lin, 2019). The AL is the primary olfactory-processing centre where olfactory sensory neurons (OSNs) from the antennae connect with projection neurons (PNs) to create globular neuropils called glomeruli. From these glomeruli, PNs connect to the MB, which are critical in olfactory learning and memory in many species (Vogt et al., 2014; Farris and Van Dyke, 2015; Groh and Rössler, 2020; Couto et al., 2023). Odours are represented in distinct glomeruli working as a functional unit, with OSNs expressing the same receptor type converging to one or two glomeruli (Zhao and McBride, 2020; Thomas, 2022). Generally, the size of a single glomerulus is assumed to reflect the number of sensory axons terminating in this structure and the size and shape of the AL, comprising all glomeruli, can be affected by odour exposure, sex and other factors (Devaud et al., 2001; Devaud et al., 2003; Giurfa, 2013; Anton et al., 2016; Grabe et al., 2016; Williams et al., 2022; Fabian and Sachse, 2023). Because of these properties, these neuropiles are good candidates to explore for plasticity in odour sensitivity at the molecular and morphological levels.

To explore molecular and morphological mechanisms of olfactory preference development we used *Bicyclus anynana* butterflies as our model species. Lepidopterans use olfaction and olfactory learning in foraging and sexual selection processes (e.g. Gowri et al. (2019); Couto et al. (2020); Dion et al. (2020); González-Rojas et al. (2020); Gowri and Monteiro (2024)), and olfactory learning has previously been demonstrated in *B. anynana*. Caterpillars can learn novel food odour preferences (Gowri et al., 2019; Gowri and Monteiro, 2024), and adult females of the species alter their male sex pheromone (MSP) preferences after a short exposure to new blends (Dion et al., 2020). For example, naive females are normally attracted to the wildtype (Wt) male sex pheromone (MSP) blend, but they can learn to prefer novel blends after a short exposure to them. The Wt MSP comprises 3 components (named MSP1, 2 and 3; (Nieberding et al., 2008; Dion et al., 2016)). Females exposed, upon emergence, to males having lower amounts of MSP1 and absence of MSP2 (called New Blend 1, NB1) lose this preference and accept mating with both NB1 and Wt males indiscriminately. Females exposed to blends with increased amounts of MSP2 (called NB2), later prefer these males over Wt males (Dion et al., 2020) (Figure 1A). The molecular details of how this odour learning process becomes encoded and retained in the brain, however, are unknown.

**Figure 1.**
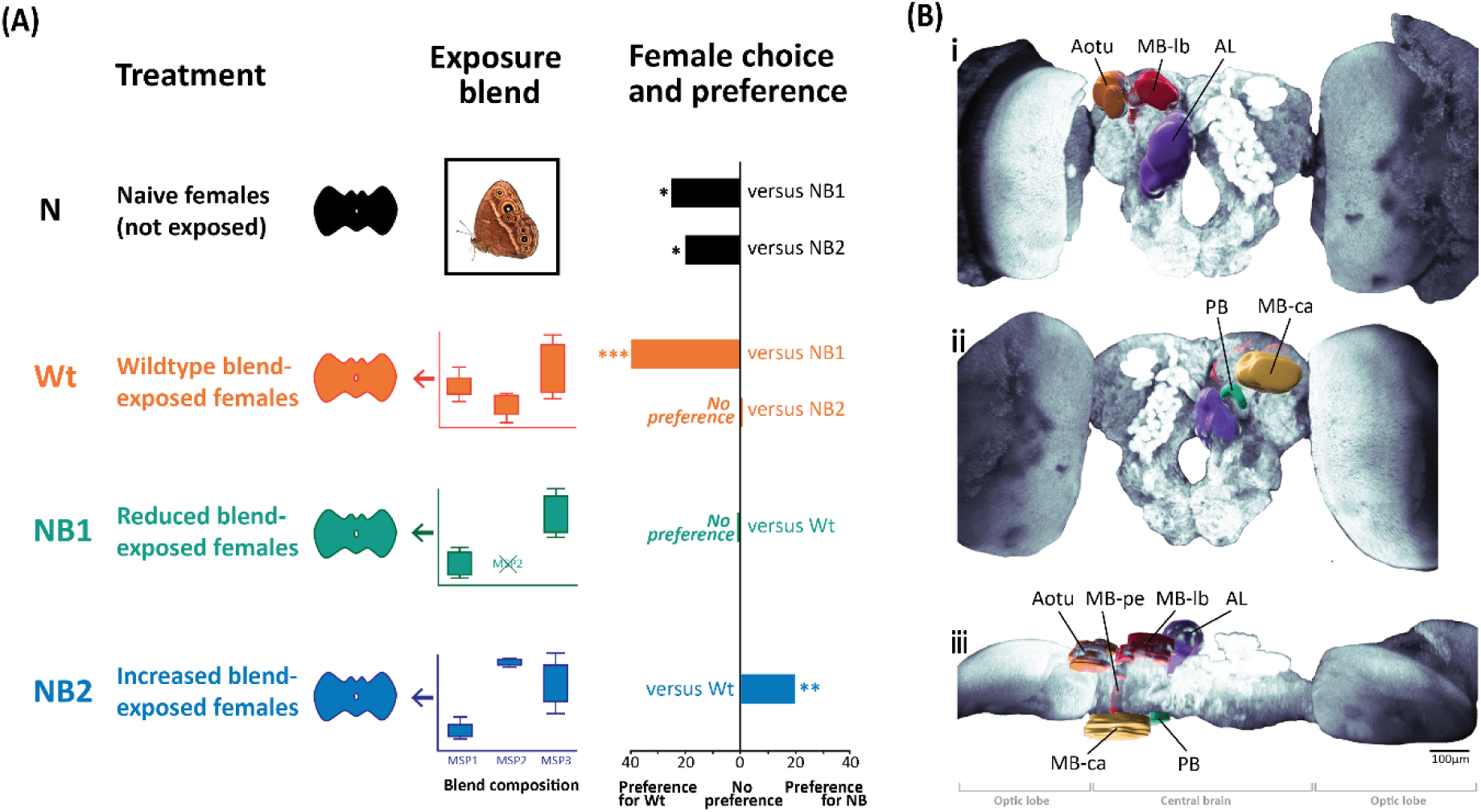
Experimental procedures and antennal lobe identification. (A) Naive females and those exposed to different pheromone blends (Treatment, left) were previously shown to have different preferences for the blends in a mate choice assay. The blend composition shows amounts of the three male sex pheromones (MSP1, 2 and 3) used for exposure (Exposure blend, middle). Naive females were isolated and not exposed until mate choice. Female preferences (right) are shown as significant increases in choice percentage from random (50%) for the Wt or new blend (NB1 and NB2) perfumed male. Asterisks illustrate significant preferences at p value <0.05 ‘*’; p<0.001 ‘**” and p< 1E-4 ‘***’ (from Dion et al, 2020). (B) 3D reconstruction of the midbrain structures shown on the animal right side from an (i) anterior, (ii) posterior and (iii) top (upper side is anterior, lower side is posterior) views. Aotu = anterior optic tubercle; AL= antennal lobe; PB = Protocerebral bridge; MB-lb = mushroom body lobe; MB-Ca=mushroom body calyx; MB-pe=mushroom body pedunculus.

Here we explore molecular and morphological correlates of pheromone olfactory learning in naïve females and in females of the same age after exposure to various pheromone blends. We also examine how male and female naïve brains differ from each other at the morphological and molecular levels. We hypothesised that brief pheromone blend exposures, as described in Dion et al. (Dion et al., 2020) (Figure 1A), would alter the volumes of specific glomeruli of the AL responding to specific MSP components, and alter gene expression and/or splicing in the female brain a few days after the exposure event. We repeated the odour exposure experiments of Dion et al. (Dion et al., 2020) (Figure 1A), dissected the brains of the butterflies on the second day (48hrs after the exposure), instead of submitting them to mate choice assays, and then either imaged the brains or subjected them to RNA sequencing. We compared naive male and female brains to identify sex-specific differences; naive versus Wt-exposed female brains to discover processes triggered by Wt MSP exposure; Wt-blend versus new blend (NB)-exposed female brains; as well as NB1 versus NB2-exposed female brains, to identify mechanisms associated with detailed perceptions of specific MSP components. We hypothesised that genes specifically related to changes in MSP2 would be up- or down-regulated in treatments with higher amounts of these components relative to other treatments (e.g., higher in NB2 versus Wt and NB1; higher in Wt versus NB1).

## Results

### *B. anynana* brain morphology is similar to that of other lepidopterans

To better visualise the brain’s finer structures, and to perform 3D volumetric reconstructions, we stained brains with an anti-synapsin antibody. The overall layout and morphology of the brain of *B. anynana* is similar to that of other Lepidoptera (*e.g.* Huetteroth and Schachtner (2005); El Jundi et al. (2009); Heinze and Reppert (2012); Montgomery et al. (2016); Sehadová et al. (2023)) (Figure 1B). The large optic lobes surround the central complex, and two pairs of antennal lobes (AL) and mushroom bodies (MB) are present on the front and the back of the central complex, respectively. *B. anynana* ALs are located at the anterior part of the brain tissue (∼10μm from the anterior of the whole brain) and span about 88μm in depth. They are distinguishable by their spherical shape on either side of the esophageal foramen, and contain "berry-like" subunits called glomeruli, arranged around the central fibrous neuropil (CFN) (Figure 2A).

**Figure 2.**
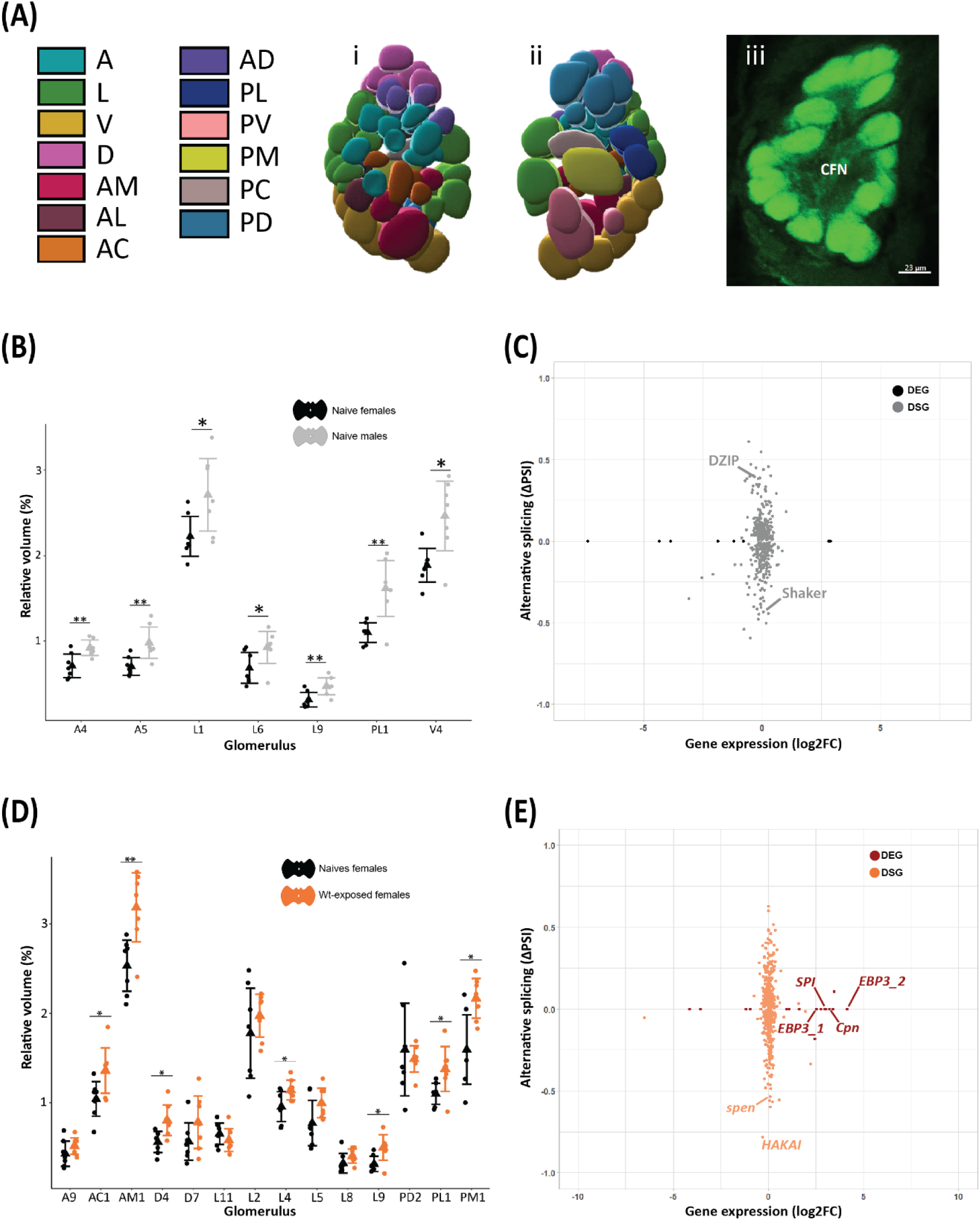
The sex of naïve butterflies and female exposure to Wt pheromone blends affect the volume of several glomeruli in the antennal lobe, and alters gene expression and splicing patterns in female brains. (A) We characterized and measured the volumes of individual *B. anynana* glomeruli in the antennal lobes. Glomeruli are colour-coded based on their spatial position within the AL, i) anterior to posterior; ii) posterior to anterior, iii) sliced image of an AL through the X-Y plane with the central fibrous neuropil labelled. A=anterior, L=lateral, V=ventral, D=dorsal, AM= Antero-Medial, AL= Antero-Lateral, AC=Antero-Central, AD=Antero-Dorsal, PL=Postero-Lateral, PV=Postero-Ventral, PM=Postero-Medial, PC=Postero-Central, PD=Postero-Dorsal, CFN=Central Fibrous Neuropil) (glomeruli positions and identification are detailed in Suppl. Fig. S2 and Suppl. File 1 Table S1); (B) Seven glomeruli were larger in naive males compared to females; (C) Magnitudes of gene expression differences (log2FC) of DEGs (padj<0.05, black dots), and inclusion level differences (ΔPSI) of DSGs (for FDR<0.05, grey dots) between naive female and male brains. (D) Seven glomeruli were significantly larger in Wt-exposed females compared to naives; (E) Magnitudes of gene expression differences of DEGs (padj<0.05), and ΔPSI of DSGs (for FDR<0.05) between naive and Wt-exposed female brains. Labelled are examples of genes involved in sensory processes or sensory neurogenesis. * above comparisons means p<0.05 and ** p<0.01, triangles are the average volumes, vertical bars are the standard errors of means, and each dot is a data point. See Suppl. File S3 for the complete list of DEGs and DSGs.

We identified an average of 67 glomeruli making up the AL in both females and males (Figure 2A, Suppl. Figure 2-1, Supp. File 2-1, Table 3), but this number varied from 64 to 69 across individuals (♀: x̄ (SD)=67±1,n=28; ♂: x̄ (SD) =67±2; n=7). The glomeruli were categorised according to the region of the AL they are located in (A, anterior; P, posterior; D, dorsal; V, ventral; L, lateral; M, medial; C, central) and assigned a unique number to differentiate them from neighbouring glomeruli in the same region (Figure 2A; Suppl.File 2-1, Table 1; Suppl. Figure 2-1). 55 glomeruli were successfully identified across all individuals (n=35) while 13 glomeruli (A2, A4, A10, AC3, AD2, AD4, AM3, D4, D6, D7, L6, L13, V10) could not be identified in at least one individual across our samples, but were still present in more than half of the individuals sampled (Suppl. File 2-1, Table 3). Glomerulus PD8 was present in all male individuals sampled, while it was only present in 18% of females.

### Seven glomeruli are larger in males than in females

To test for sex-specific differences in the AL, we compared the sizes of glomeruli of naive males and females. Both sexes had similar total AL volume (**♀:** x̄=1.14×10^6^μm^3^, **♂:** x̄=1.25×10^6^μm^3^ ; t_10_=0.556, P=0.59) and similar total volume of all glomeruli (**♀:** x̄=7.76×10^6^μm^3^, **♂:** x̄=9.23×10^6^μm^3^ ; t_11_=0.432, P=0.674) (Suppl. File 2-1, Table 2). The 10 largest glomeruli for both females and males were similar in volume, with some slight differences in ranking (Suppl. File 2-1, Table 4). Seven glomeruli (A4, A5, L1, L6, L9, PL1 and V4), positioned at different locations in the AL, were significantly larger in males than in females, while all other glomeruli had similar sizes across the sexes (Figure 2B).

### Exposure to Wt and new pheromone blends affected the size of specific glomeruli in female brains

To test whether exposure to Wt or new pheromone blends altered the size of glomeruli, we compared the volume of glomeruli of naive and exposed females after immunostaining. In Wt-blend exposed brains, seven glomeruli were significantly larger relative to naive female brains (AC1, AM1, D4, L4, L9, PL1 and PM1), with two glomeruli from this list being larger in naive males compared to naive females (L9 and PL1) (Figure 2D; Suppl. File 2-1 Table 5). In females exposed to NB1 blend (reduced blend) two glomeruli were smaller compared to those exposed to Wt blend (D7 and PD2), while two other glomeruli, L11 and L8, were larger (Figure 3A; Supp. File 2-1 Table 5). These four glomeruli had similar sizes between naive and Wt-exposed females. In NB2-exposed females (increased blend), two glomeruli were larger (A9 and L2) and two were smaller (D4 and L5) than in Wt-exposed females (Figure 3C; Suppl. File 2-1 Table 5). All glomeruli had a similar size between NB1- and NB2-exposed brains (Figure 3E, Supp. File 2-1 Table 5). There was no glomerulus that commonly shared a change in size across the different comparisons.

**Figure 3.**
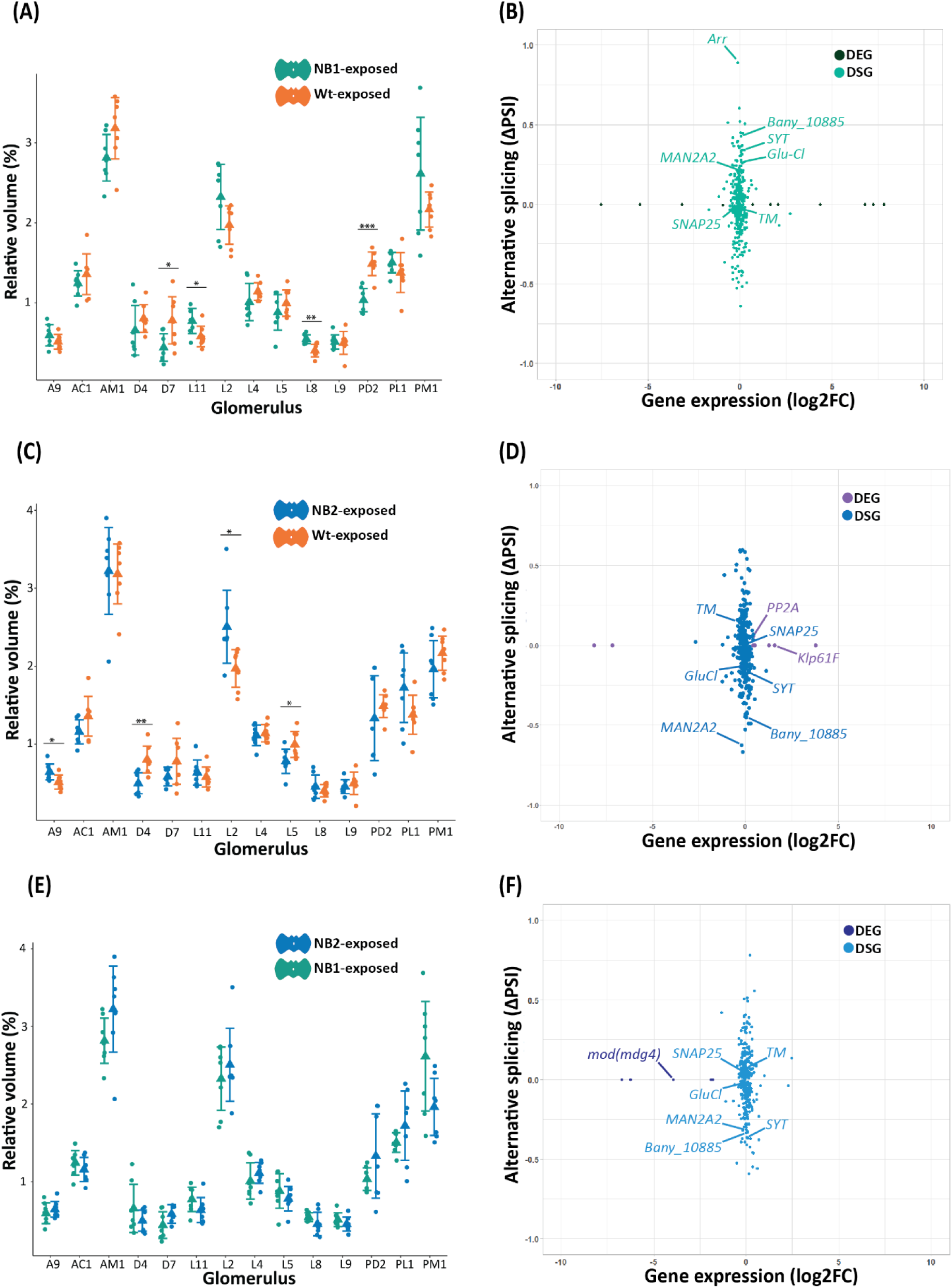
Exposure to new pheromone blends affected the volume of specific glomeruli, and gene expression and splicing patterns in female brains. (A), (C) and (E) are glomeruli size differences observed between Wt and NB1, Wt and NB2, and NB1 and NB2 exposed female brains respectively (with * above comparison for p<0.05, ** for p<0.01, *** for p<0.001, triangle is the average volume, vertical bar the standard error of the mean, and each dot is a data point). (B), (D) and (F) show the gene expression differences (log2FC) of DEGs (padj<0.05), and inclusion level differences (ΔPSI) of DSGs (for FDR<0.05) in the Wt versus NB1, Wt versus NB2, and NB1 versus NB2 comparisons. Labelled are examples of genes involved in neurogenesis, metabolism and development.

### The number of OR genes, expressed at low levels in the brain, approximates the number of glomeruli

Because the number of glomeruli closely resembles the number or ORs normally expressed in cells of the peripheral nervous system, such as the antennal tips (Carlsson et al., 2013), we examined the number of transcripts for chemical receptors and olfactory-related proteins that are expressed in the brain using our transcriptome sequences. We counted 55 olfactory receptors (OR) genes, the OR co-receptor ORCO, and 3 joint olfactory and gustatory receptors (OGR). In the transcriptome, we also found 41 additional gustatory receptors (GR), 14 olfactory binding proteins (OBP) including one pheromone-specific (PBP), 41 ionotropic receptors (IR), 6 sensory neuron membrane proteins (SNMP), one chemosensory protein (CSP) and 20 members of the ejaculatory bulb protein 3 (EBP3) family, also known as chemosensory protein homologs, as pherokine 3, or as the odorant-binding protein A10 (McKenna et al., 1994; Pikielny et al., 1994; Bohbot et al., 1998). We also identified 10 pickpocket (*ppk*) proteins and 5 transient receptor potential channels (*trp*) (Supplementary File 2, Table S5). Most of these proteins (about 70%) are expressed at very low levels in the brains, with less than 10 cumulative read counts over all libraries. It is interesting to note, however, that we identified 55 ORs and that only 55 glomeruli were consistently identified across all individuals, suggesting a close association or specificity between ORs and glomeruli number.

### Sex pheromone exposure affects gene splicing more than gene expression

We generated RNAseq libraries from the butterfly brains to evaluate the impact of sex and pheromone blend exposure on gene expression and splicing. The PCA done on a random sample of 5000 gene counts and the hierarchical clustering analysis showed low levels of clustering according to treatment or sex, with a small variance across samples (cumulative 38%), suggesting that brain exposure to pheromones and butterfly sex don’t correlate with large shifts in gene expression (Supplementary Figure S3A and S3B). Three outliers belonging to naive males, NB1-exposed and the NB2-exposed females deviated significantly from the cluster on the PCA and were removed from the dataset before further differential expression analysis. In all comparisons, the differentially expressed genes (DEGs) were a different set of genes than the differentially spliced genes (DSGs). No gene was both DE and DS (Figure 2B, 3G, 3H, 3I; Supplementary Figure S4B). A small number of 18 to 32 genes were identified as DEGs (Padj < 0.05) between treatments. The highest number of DEGs were found in the naive versus Wt-exposed comparisons (Supplementary File 2, Table S2; Supplementary File 3). No enrichment was found in any of the DEG clusters based on gene ontology. In contrast, an average of 55,470 alternative splicing events were detected in a total of 1937 genes across libraries, including an average of 857 DS variants across all comparisons (FDR<0.05, IJC and SJC >20), with most differences also being found between the naive versus Wt-exposed comparison (1048 significant splice variants). Overall, a higher proportion of retained intron sites (∼31% on average), followed by spliced exon sites (average ∼26%), were detected relative to the other types of splicing events (Supplementary File S2, Table S3; Figure S3C & S3D).

### Naive male and female brains have similar gene expression but different splicing patterns

To examine sex-specific gene expression and splicing in *B. anynana* brains, we compared data from naïve male and female brains. We found a small number of 9 downregulated and 9 upregulated genes in female brains compared to males, but a much larger number, and distinct set of 394 genes, having significantly different splicing patterns between the sexes (Supplementary Figure S3C, S4C). Among the DEGs, 13 were uncharacterized or non-annotated proteins. The remaining identified DEGs were mostly involved in membrane building and structural constitution of the cuticle, or cellular signalling and homeostasis (Supplementary File 3). The GO analysis revealed that the DSGs were primarily involved in calcium binding and organ morphogenesis, mostly at the cell periphery and cell projections (Supplementary Figure S3E; File S2 Table S4). The potassium voltage-gated channel *Shaker* and the zinc finger protein DZIP1 are examples of such genes with high significant absolute PSI (0.43 and 0.35 respectively). Interestingly, genes from the sex-determination pathway, such as *doublesex*, *transformer*, *sex-lethal* and *fruitless* didn’t show sex-biased expression of alternative transcripts in the naive butterfly brains.

### Exposure to Wt males induces the upregulation of several sensory-related genes in female brains

To test whether females exposed to Wt male sex pheromones changed their gene expression and splicing patterns relative to naive females of the same age, we performed DEGs and DSGs analyses (Supplementary File S2 Table S2). Compared to naives, 14 genes were downregulated, and 16 genes upregulated, in Wt-exposed females. The upregulated set contained sensory-related genes such as both copies of the ejaculatory bulb-specific protein 3 (EBP3), also known as chemosensory protein (CSP) or insect odorant-binding protein A10 (Pikielny et al., 1994; Angeli et al., 1999), the olfactory neuron axon outgrowth protein alaserpin (SPI, serine protease inhibitor), and the eye-specific gene *calphotin* (*Cpn*), all between 2.5 and 4 times upregulated in exposed females (Figure 2B, Supplementary File S3). Exposure to the Wt blend induced 474 significant DS events (at p<0.05), with an overrepresentation of genes involved in RNA binding, microtubule binding and mitochondrial inner membrane functions (Figure S3E; Supplementary File S2 Table S4). An example of a DSG of a binding protein, with some of the largest values of inclusion level differences between the two treatments, include the ubiquitin-protein ligase Hakai, affecting RNA splicing and methylation (A5SS, ΔPSI=-0.78, fdr=1.6749e-10; Figure 4C) in fruit flies.

**Figure 4.**
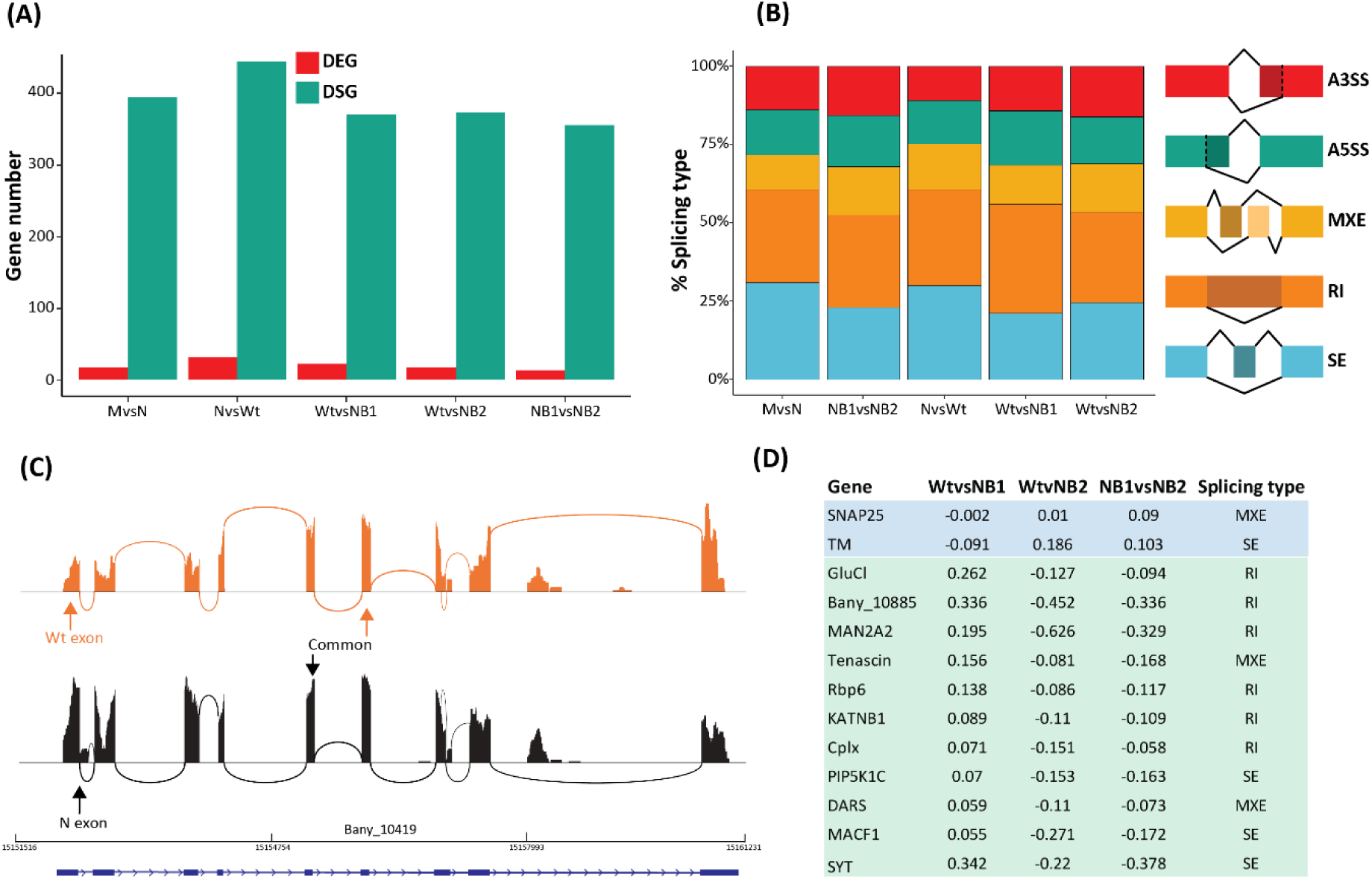
There are more significant changes in splicing events than changes in gene expression between treatments. (A) number of DEG and DSG identified in the different comparisons; (B) proportions of significant splicing types found in the different comparisons (at FDR<0.05, IJC and SJC >20); (C) PSI of genes associated with an increase and a decrease in MSP2 levels (in blue and green, respectively); (D) sashimi plots showing HAKAI A5SS isoforms in a Wt and Naïve sample.

### Exposure to new blends affects the expression of cellular growth and developmental genes, and the splicing pattern of RNA binding genes

To identify genes affected by the sex pheromone components, we identified those commonly down- and up-regulated in the Wt versus NB comparisons, and between NB1 and NB2. Twenty-three genes were DE in the Wt vs NB1 comparison (30% were not annotated or characterised (NAC)), nineteen were DE in the Wt vs NB2 comparison (37% NAC), and fifteen genes were DE in the NB1 vs NB2 comparison (67% NAC) (Supplementary File S3). The annotated DEGs were mostly involved in DNA biosynthetic processes and catalytic activities. For instance, the cell growth control and development protein phosphatase *PP2A* and the cell proliferation kinesin-like protein *Klp61F* were both upregulated in NB2-exposed brains compared to Wt, (Figure 3D) while a chromatin remodelling *mod(mdg4)-like* gene was downregulated in NB2 compared to NB1 exposed brains (Figure 3F). There is no gene that is both upregulated in NB2 (versus Wt, and versus NB1) and downregulated in NB1 (versus Wt). Similarly, no gene is downregulated in NB2 (versus Wt and NB1) and upregulated in NB1 (versus Wt).

Most of the differences observed across treatments were in differential gene splicing. DSGs involved in mitochondrial inner membrane and RNA binding were over-represented in the Wt versus NB1 and in NB1 versus NB2 comparisons. For example, the gene with the highest value of inclusion level difference (PSI of 89%) in NB1 (compared to Wt) is the regulator of GPCR (G Protein Coupled receptor) signal transduction arrestin (*Arr*), also having different splice variants in the other treatment groups (Figure 3B). Other DSGs with high inclusion, and found in the three groups, function in neurogenesis (e.g. the neurite growth and transcription factor Zinc-finger protein *DZIP* (Bu et al., 2021). To identify splicing events specifically affected by MSP2 levels, we selected variants that have a higher value of inclusion level difference in NB2 compared to Wt and NB1, and in Wt compared to NB1. We found 2 genes annotated as synaptosomal-Associated Protein 25 (SNAP25) and tropomyosin (TM). The percent splice indexes for these genes are lower than 10% in most comparisons (Figure 4C; Suppl. File 2, Table S6). To connect splicing events to exposure to low MSP2 levels, we identified 11 genes with variants that have a lower percent splice in index in NB2 compared to Wt and to NB1, and in Wt compared to NB1. Among these, synaptotagmin (SYT), alpha-mannosidase (MAN2A2), a glutamate-gated chloride channel (GluCl) and an uncharacterized ion transporter (Bany_10885) have the highest absolute PSI in the treatment comparisons (above 20%) (Figures 3B; 3D; 3F; 4D, Suppl. File 2 and Table S6).

## Discussion

In previous work we showed that a brief, three-minute exposure of naive female butterflies to male sex pheromone blends, that differ in the quantity of at least one or two known pheromone components, led to changes in female preferences for the male blends, two days after the exposure treatment. Here we show that these brief pheromone exposures affected the size of a subset of antennal lobe glomeruli, altered the expression level of a handful of genes, and affected the splicing of hundreds of genes in the brains of females. We also show that antennal lobe morphology is sexually dimorphic, and that male and female brains do not differ much in gene expression but are massively different in gene splicing patterns.

### Some glomeruli are larger in males and several glomeruli show size plasticity in response to odour experience

Naive male and female brains have similar morphologies upon emergence. The number of glomeruli, total AL volume, and the volume of all glomeruli were similar in the two sexes, which is consistent with previous studies in the Monarch *Danaus plexippus* and the Neotropical butterfly *Godyris zavaleta*. These studies have identified 65-70 and 67 glomeruli on average respectively in the two species, with no sex specific differences in total AL volume (Heinze and Reppert, 2012; Montgomery and Ott, 2015). Naive *B. anynana* males had, however, seven glomeruli larger than females, and one of them became larger in female brains after they were exposed to Wt blends. These sexually dimorphic glomeruli are possibly responding to still unknown olfactory cues used primarily by males of the species. They are likely not involved in sex pheromone perception (Costanzo and Monteiro, 2007; Nieberding et al., 2008), as six of these glomeruli didn’t change in size in the exposed females. It is also possible that MSP-specific glomeruli are not sexually dimorphic if males can detect MSP from other males while in competition for a mate. The pattern of responses of Lepidopteran glomeruli to volatiles is complex, however, with different glomeruli being able to respond to the same compound, and the same glomeruli being able to react to different compounds (e.g. (Trona et al., 2010; Varela et al., 2011; Larsdotter-Mellström et al., 2016; Herre et al., 2022)). The functional significance of sexually dimorphic glomeruli in *B. anynana* remains an open question.

Changes in glomeruli volumes following the exposure events were most likely due to cellular growth or differentiation of innervating OSNs, and changes in the number of synaptic connections they form with local interneurons and projection neurons (Grabe et al, 2016, Fabian, 2023). This is because the DEGs and DSGs identified were involved in gene expression regulation (RNA binding, transcription factors) and neuronal tissue development and maturation. For instance, DEG *Klp61F* impacts axons and dendrite microtubule growth (Heck et al., 1993; Feng et al., 2021), and DEG *mod(mdg4)* affects synaptic plasticity in the *Drosophila* brain (Lin et al., 2021). DSG *arrestin* regulates the signalling of G coupled receptors, impacting the development of the olfactory transduction machinery. An arrestin-related protein called mKast is also expressed in pupal and adult honeybee kenyon cells, neurons of the MB. It is thought to affect cell fate but its exact function is still unknown (Raming et al., 1993; Alvarez, 2008; Kaneko et al., 2016; Yamane et al., 2017; Kohno and Kubo, 2018). In *B. anynana*, the short three-minute exposure to the male blends was enough to create cellular growth or differentiation of new neurons in the females’ sensory and memory structures two days later, but the precise mechanism behind this glomerular plasticity needs to be determined.

### Naïve brains were sexual dimorphic primarily in gene splicing

Naive brains exhibited few sex-specific differences in gene expression, with the few DEGs identified being involved in cuticle development. Sex differences in gene expression in *D*. *melanogaster* brains were also previously shown to be low, with male heads being closer to female heads than to other male tissues (Catalán et al., 2012; Gibilisco et al., 2016). In insects, sex-biased gene expression was found in reproductive tissues (e.g. *in Drosophila* (Parisi et al., 2003; Gibilisco et al., 2016)); while in heads, sex-biased gene expression occurs primarily in non-nervous tissue (Goldman and Arbeitman, 2007; Chang et al., 2011). Cuticle genes being primarily DE in the male and female brains of *B. anynana* supports these findings.

Alternative splicing, however, appears to be an important mechanism governing sex differences in the brains of young butterflies. In both invertebrate and vertebrate species, such as fruit flies, fishes, bats and humans, the sexes show large differences in gene splicing (Wang et al., 2008; Trabzuni et al., 2013; Mohr and Hartmann, 2014; Gibilisco et al., 2016; Karlebach et al., 2020; Naftaly et al., 2021; Chen et al., 2023), and brains/heads usually show some of the highest transcriptional diversity (along with gonads) across all tissues. Generally, sex-specific spliced genes are thought to derive from the sex differentiation pathway, crucial to neurological development. In insects, these genes include the sex-specific isoforms of *doublesex* (*dsx*) and *fruitless* (*fru*) (Prakash and Monteiro, 2016; Hopkins and Kopp, 2021), which are known to impact sexual behaviours and sex-specific morphologies across species (Salvemini et al., 2010; Wexler et al., 2019; Prakash and Monteiro, 2020; Takahashi et al., 2021). In our study it was surprising that none of the well-known sex differentiation pathway genes were DS (or DE) between naive brains, unlike hundreds of other genes. It is possible that sex-specific variants of these genes might be affecting brains at earlier stages of development, as it is the case for *fru* in *Drosophila* larval nervous system development (Neville et al., 2014). Alternatively, the recent discovery that *B. anynana* uses a unique primary sex determination signal (Van’t Hof et al., 2024), distinct from that used in *Drosophila* or *Bombyx mori*, could have led to yet unexplored diversification of the sex-determination pathway in this species.

Gene ontology analysis for the set of DSGs indicated that male and female brains differed in signalling processes involved in neurogenesis and happening mostly at axons or dendrites. Calcium ion binding proteins, which function in signal transduction and regulate many aspects of the cell’s functioning, homeostasis, neuron excitation, and synaptic transmission were overrepresented in the set of DSGs (Luan and Wang, 2021; Saghian and Wang, 2022). DSGs involved in brain morphogenesis, and located at the cell projection and periphery were also overrepresented. An interesting candidate DSG with a high PSI was *Shaker* (*Sh*), a potassium voltage-gated channel known to be expressed as multiple splice variants that encode proteins with distinct structural features in the *Drosophila* brain (Kamb et al., 1988; Pongs et al., 1988; Schwarz et al., 1988). The response of cation channel splice variants to the same ligand can be very different, *e.g.* it can lead to opposite responses to an insecticide (Tan et al., 2002), but the functions of *Sh* variants are unknown. *Drosophila* brain *Sh* expression affects male sex discrimination (Houot et al., 2012), visual motion detection, and photoreceptor sensitivity (Peretz et al., 1998; Gür et al., 2020), so the different variants expressed in male and female *anynana* brains should be examined in future in connection to similar behavioural essays.

### Changes in glomeruli volume do not correlate with female exposure to specific MSP blends

There is little support for the role of altered glomeruli volume in female preference change for a new sex pheromone blend. Changes in specific MSP did not lead to consistent and specific glomeruli size changes, and thus we could not identify the glomeruli tuned to the specific pheromone components. MSP quantity and ratio changes between Wt, NB1, and NB2 males might be too minor to impact glomeruli volume, or our sampling time, 2 days after the exposure, might have been too soon to detect size differences. Alternatively, odour exposure alters female behaviour via molecular mechanisms that do not change brain morphology.

### Expression differences in an odorant binding protein may be associated with MSP3 learning

The OBP homolog EBP3, an odorant binding protein (OBP), could be involved in the butterfly olfactory perception and learning process of MSP3. Two EBP3 copies were upregulated in the brains of females exposed to Wt blends relative to naive individuals. Because these two *EBP3* genes were not differentially expressed in the other comparisons, these OBPs could be binding to MSP3 specifically, as this pheromone is present in similar amounts in Wt, NB1 and NB2 males (Figure 1A). EBP3 carries semiochemicals with different proposed roles, and is also known as odorant-binding protein A10 (OBP A10), P10, CSP or pherokine, among other names (McKenna et al., 1994; Pikielny et al., 1994; Bohbot et al., 1998; Angeli et al., 1999; Jacquin-Joly et al., 2001; Sabatier et al., 2003; Pelosi et al., 2018). OBP A10 is known to bind to several pheromone components from fruit flies and moths (e.g. Bohbot et al. (1998); Kitabayashi et al. (1998); Nagnan-Le Meillour et al. (2000); Jacquin-Joly et al. (2001)), and to react to plant volatiles in pest flies (Zhu et al., 2023) and psyllids, where its upregulation is correlated with increased preference for phytochemicals (Lin et al., 2022). In *D. melanogaster*, a mushroom body-expressed *EBP3* was recently associated with long term memory formation (Widmer et al., 2018), and other OBPs were previously shown to be differentially regulated in the brains of bees after olfactory learning (Wang et al., 2013). Here, the two *EBP3s* might be relevant candidates for blend preference memory formation in exposed females. Functional analyses for these genes are needed in *B. anynana*, with perhaps a focus on mushroom bodies, to elucidate their role in memory formation.

Apart from *EBP3*, there is little support for changes in gene expression correlating with changes in female preference for new sex pheromone blends. MSP quantity and ratio changes between Wt, NB1, and NB2 males might be too minor to impact gene expression, or it may impact gene expression at different time points. Most importantly, small gene expression changes in the mushroom bodies, the main centres for learning in insects (Zars, 2000; Giurfa, 2013; Falibene et al., 2015; Baltruschat et al., 2021), might not have been detected in our whole-brain transcriptomics analysis and may require a single-cell sequencing approach.

There were, however, a few other differences in gene expression between naive and Wt-exposed females that might have affected olfactory perception and growth of sensory structures. For example, two Wt-female up-regulated genes, *calphotin* and *alaserpin* (SERA or antichymotrypsin-2, a serine protease inhibitor) indicate the growth of visual and/or olfactory organs. *Calphotin* functions in rhabdomere development (Martin et al., 1993; Charlton-Perkins and Cook, 2010; Allen et al., 2018) and *alaserpin* functions in antennal neuron extension (Hanneman and Kanost, 1992). The new sensory experience of these Wt pheromone-exposed females may have promoted the expansion of associated sensory organs needed by females to navigate their new environment.

### Gene splicing is different between pheromone exposure treatments

A much larger number of genes were differentially spliced, relative to differentially expressed, in response to early exposure to new sex pheromone blends. In addition, genes were exclusively differentially spliced or up- and down-regulated in the different treatments. Similar results, of non-overlapping gene sets either DS or DE, have been described in aphid polyphenic morphs or in butterfly wings from different seasonal forms, including *B. anynana* (Kalsotra and Cooper, 2011; Grantham and Brisson, 2018; Tian and Monteiro, 2022).

We found that DSGs with RNA and microtubule binding functions were overrepresented in the Naive vs Wt, and NB1 vs NB2 comparisons. This suggests a role for the splice variants in the formation of neuronal networks that involve substantial RNA processing and cytoskeletal modifications. *Drosophila* and mammalian nervous systems are also known to use alternative splicing at high frequency for mechanisms like cell differentiation and morphogenesis (Norris and Calarco, 2012; Zheng and Black, 2013; Mohr and Hartmann, 2014; Su et al., 2018).

We noted a higher proportion of IR events, where unspliced introns remain in mature mRNA, relative to other types of splicing events. IRs were shown to affect RNA binding proteins and contribute to the plasticity of the transcriptome and regulation of gene expression during cell development, cell differentiation, and in response to cellular stress (Grabski et al., 2021; Wong and Schmitz, 2022). In mammals, IR affects the synaptic plasticity of neuronal cells, and can be triggered by a short stimulation, allowing a fast neuronal plastic response to a stimulus (Buckley et al., 2011; Mauger et al., 2016; Petrić Howe et al., 2022). The precise role of the IR variants and how they affect pheromone perception and learning in butterflies remains to be investigated.

A learned preference for new blends was linked to variants of proteins involved in synaptic transmission. Levels of expression of these proteins affect learning and memory in other species. For instance, synaptic Glutamate-gated chloride channels regulate fast inhibitory synaptic transmission, and have 6 known variants (across insects) with different spatial and developmental expression patterns, and response strength to the substrates (neurotransmitter and insecticides) (Furutani et al., 2014; Kita et al., 2014; Wu et al., 2017). These channel proteins are necessary for plant volatile olfactory memory retention in honeybees (El Hassani et al., 2012; Démares et al., 2014). Synaptotagmin and SNAP25 interact to promote neurotransmission and synaptic growth in *Drosophila* (Schiavo et al., 1997; Kikuma et al., 2017). SYT are calcium sensors with multiple variants (Dean et al., 2012), where the syt-1 variant affects long-term synapse potentiation and promotes learning in mice (Wu et al., 2020). SNAP25 is part of the attachment protein receptor complex responsible for exocytosis at synapses in insects (Johard et al., 1999; Vilinsky et al., 2002; Nagy et al., 2005). Bee olfactory learning reduces SNAP25 expression levels (Zhang et al., 2015), and the protein is involved in memory consolidation in rats (Ren et al., 2018) and humans (Hou et al., 2004; Spellmann et al., 2008; Golimbet et al., 2010). However, the roles of specific variants of these proteins in butterflies remain to be tested.

In summary, we showed extensive sexual dimorphism in gene splicing in male and female *B. anynana* butterfly brains, and showed that a short exposure of a female butterfly brain to various male pheromone blends alters the size of a few glomeruli in the antennal lobe and changes additional splicing patterns in the brain. Specific changes in one pheromone component particularly affected the splicing of genes involved in synaptic transmission. Functional experiments will need to be performed in some of the genes identified in this study to probe how and which of these changes mediate pheromone odour learning that ultimately impacts female mate choice.

## Materials and Methods

### Insect rearing and odour treatments

Insect production and odour treatments were the same as in (Dion et al., 2020). Individuals were reared at 27°C, 80% humidity and 12:12 h light:dark photoperiod. Caterpillars and butterflies were fed unlimited corn leaves and mash banana, respectively. Sexes were separated at the pupal stage, with each pupa placed individually in a plastic box and, upon emergence, in plastic cages. Males used for expose to females were kept in groups in age-specific cages. After blend manipulation, males were allowed to rest for 30 minutes, and then manually exposed to the females. Males were all aged 4 to 6 days old and all animals were virgins. Females within 3 hours of emergence were exposed to males for 3 minutes, and then isolated in cups for 2 days (∼48 hrs). This protocol was used previously to identify changes in female behaviour. The treatments (male sex pheromone blends used for female exposure), and female responses were as follows (Figure 1A):

● Naive females (N) and naïve males (M) were kept isolated in individual boxes until dissection on day 2. They were not fed or used for behavioural experiments. Naive females have significant preferences for the Wt blend when tested against NB1 or NB2 in choice assays.
● Wt-exposed females (Wt) were manually exposed upon emergence to males carrying the Wt blend, then they were placed in individual containers and dissected two days later. These females have a strong preference for the Wt males when tested against NB1 males, but they randomly choose between Wt and NB2 males.
● NB1- and NB2-exposed females (NB1 and NB2) were manually exposed upon emergence to males carrying the respective reduced and increased blends, placed in individual containers, and dissected two days later. NB1-exposed females choose randomly between the two males. NB2-exposed females significantly prefer NB2 males.

### Immunohistochemistry

Brains were dissected, fixed, and stained following prior protocols (Ott, 2008; Toh et al., 2021) with minor modifications. Briefly, the head was submerged in HEPES-buffered saline (HBS; 150mM NaCl; 5mM KCl; 5mM CaCl2; 25mM sucrose; 10mM HEPES; pH 7.4) and fixed for 22-24 hours under agitation at room temperature in zinc-formaldehyde solution (ZnFa; 0.25% [18.4mM] ZnCl2; 0.788% [135mM] NaCl; 1.2% [35mM] sucrose; 4% formaldehyde). The brains were then dissected in HBS and washed 3 times in HBS before further staining. Samples were then submerged in 80% methanol/ 20% dimethyl sulfoxide (DMSO) for 2 hours under agitation and subsequently stored in 100% methanol in -20°C till further processing. To prepare these samples for staining, they were brought to room temperature and rehydrated in decreasing methanol concentrations (90%, 70%, 50%, 30%, 0% in 0.1M Tris buffer, pH 7.4, 10 minutes each). Subsequently, they were pre-incubated for 1.5 hour at room temperature in a mixture of 5% Bovine Serum Albumin (BSA; Sigma Aldrich, St. Louis, Missouri, United States) and 0.1 M phosphate-buffered saline (PBS; pH 7.4 containing 1% DMSO and 0.005% NaN3) (PBSd). They were then stained with synapsin antibodies (anti-SYNORF1 obtained from the Developmental Studies Hybridoma Bank DSHB, product 3C11; https://dshb.biology.uiowa.edu/) at a 1:30 dilution in PBSd-BSA for 3.5 days at 4°C and washed thereafter with PBSd (3x 1.5 hours). Alexa Fluor 488 (AF 488) conjugated goat anti-mouse IgG antibody (Thermofisher Scientific) was added in a 1:100 PBSd-BSA ratio for 2.5 days at 4°C. The brains were then dehydrated in increasing glycerol concentrations (4% for 2 hours, 15%, 50% and 80% for 1 hour each) in 0.1M Tris buffer with DMSO to 1%. After which they were submerged into 100% ethanol in a drop of 80% glycerol. The ethanol was agitated for 20 mins and refreshed twice (30 mins each) without agitation. Clearing of the brain tissues was done by dripping methyl salicylate into brains immersed in 100% ethanol and waiting for the brains to sink before refreshing the methyl salicylate once (30 minutes incubations).

### Confocal Imaging

The cleared brains were mounted in methyl salicylate onto glass slides with a depression of 0.6-0.8mm (Marienfeld Superior, Germany) and the coverslip sealed with clear nail polish. All imaging was done using the Olympus FV3000 confocal laser-scanning microscope with either the 4x or 10x dry objective lens and an aperture of 120μm. A 488-nm laser line was used to excite the dye (AF 488). For each whole brain sample, we visually identified the different brain structures (Figure 1 B) and took a stack of images from either the left or right Antennal Lobe (AL) selected at random using the 10x air objective with the following image settings: a mechanical z-step of 2 μm and an x-y resolution of 1024 × 1024 pixels. An additional zoom factor of 4.21x was applied for optimal z sampling. Finally, a 1.52x correction factor was applied to the voxel size in the z-dimension to correct for the artifactual shortening caused by the 10x air objective (Heinze and Reppert, 2012).

Imaging for the whole brain required stacking two stitched side by side images taken with an overlap of 10%. We used the 4x objective with a mechanical z-step of 2 μm and an x-y resolution of 512 x 512 pixels. Stacking and stitching were done in FIJI (Schindelin et al., 2012).

### Glomerular identification and volumetric reconstructions

We first identified each glomerulus based on their location, size and shape within each AL sample by comparing the confocal stacks directly at different depths throughout the AL. Several easily distinguishable glomeruli were used as landmarks paving the way for identification of the other glomeruli in the region (Figure 2A; Suppl. File 1, Table S1; Suppl. Figure 2-1). We followed the nomenclature used in several other studies including Solari et al. (2016) and Stocker et al. (1990), for naming of the glomerulus in *B. anynana*.

After identification, each glomerulus was reconstructed in Imaris 9.9.0 (Oxford Instruments Group) based on the stack of confocal images taken. We manually delineated individual glomerulus using the Imaris *Add New Surface* module by outlining with the *Draw* function the boundaries of each. We performed the outline on every alternate image slice within one stack for each respective glomerulus. Then a surface rendering of the glomeruli was done with the *Create Surface* function to obtain a 3D model, allowing us to visualise the spatial position of each glomerulus within the antennal lobe. The colours of the glomeruli were assigned using the *Colour* tab and glomeruli belonging to the same region were coded in the same colour (Figure 2A; Suppl. Figure 2-1; Supplementary File 2-1, Table1). We subsequently obtained the volume of each glomerulus using the *Statistic* tab. We also obtained, in a similar fashion, the volume of the whole AL (consisting of the volume of all glomeruli subunits and the central fibrous neuropil (Figure 2A; Supplementary File 2-1, Table 2). A reference key detailing the process of glomerulus identification based on the relative positions to the landmark glomeruli is available in Suupl. File 2-1, Table 1; Suppl. Figure 2-1 and Suppl. Results 2-1.

### Glomeruli volume statistical analysis

All comparisons were performed with a two-tailed Student’s t test unless otherwise stated. To account for the individual variation in body size of each sample, we normalised a) the absolute volume of the whole AL to the volume of the midbrain for comparisons of total AL volume and b) the volume of each glomeruli to total glomerular volume (total AL volume excluding the CFN; Figure 2A) for comparisons between naive males and females. We also normalised the absolute volume of each glomerulus to the volume of the whole AL for comparisons between females exposed to different MSP blends. If the relative volumes were found to deviate from normality through the Shapiro Wilk’s test, non-parametric Mann-Whitney U test was used instead for comparison. All statistical analyses were done using R Statistical Software v4.0.1 (R Development Core Team, 2020) and graphs were plotted with the plyr v1.8.8 (Wickham, 2011) and the ggplot2 v3.3.6 (Wickham, 2016) packages.

### RNA extraction, sequencing, and assembly building

Two days after exposure and isolation, butterflies were flash frozen in liquid nitrogen, their brains dissected following Toh et al. (2021) and stored in RNALater (Invitrogen, ThermoFischer Scientific, no. 7020). Five brains were pooled per replicate, and five replicates were made for each of the five treatments. RNA extraction was performed using phenol and chloroform. The samples and RNA extractions were randomised to minimise batch effects. The 25 samples were sent to NovogeneAIT Genomics, Singapore, for library preparation and RNA sequencing.

The initial quality control of the sequencing files was performed with FastQC v.0.11.9 (https://www.bioinformatics.babraham.ac.uk/projects/fastqc), followed by adaptor trimming and cleaning of low-quality raw reads with Trimmomatics v0.39 (Bolger et al., 2014). The bbsplit script from the bbmap toolbox (Bushnell, 2014) was used to filter-out the bacterial sequences from the dataset by aligning the reads to a concatenation of bacterial genomes downloaded from the National Center for Biotechnology Information (NCBI) in June 2018. We identified and removed ribosomal RNA sequences by aligning the reads to the eukaryotic rRNA database available in sortmeRNA (Kopylova et al., 2012). We obtained the gene and transcript count matrices by aligning the clean reads to the *B. anynana* genome V2 (Murugesan et al., 2022) in Hisat2 (Pertea et al., 2016; Kim et al., 2019). Resulting .bam files for each library were aligned to the .gff3 *B. anynana* genome with stringtie and the abundance tables were compiled with the python prepDE.py script (Pertea et al., 2016). On average, 86.49% of clean reads mapped to the genome (Suppl. File 3-1, Table 1). For local blast purposes, we used stringtie to build a brain assembly by converting the libraries’ .bam to .gtf files that were merged into the assembly file and converted to a fasta file with the gffread utility (Pertea et al., 2015; Pertea et al., 2016; Pertea and Pertea, 2020).

### Gene expression analysis

Principal component analysis (PCA) plots were built on normalised gene counts using ggplot2 (Wickham, 2016) in R (R Development Core Team, 2020). Three libraries (one M, one NB1 and one NB2) were excluded from the DE analysis as they were considered outliers based on the PCA (Suppl. Figure 3-1). We identified differentially expressed genes (DEGs) from the gene count matrix using the DESeq2 package (Love et al., 2014) in R v4.1.3 (R Development Core Team, 2020) implemented in RStudio V2022.07.01 (Posit Team, 2023). DEGs were identified using the ‘contrast’ command to perform the comparisons between naive males and naive females, naive females and Wt-exposed females, Wt-exposed females and NB1-exposed females, Wt-exposed females and NB2-exposed female, and finally NB1- and NB2-exposed females. All genes with less than 10 reads in total over all libraries were filtered out. Log fold changes were transformed using ‘apeglm’ (Zhu et al., 2018) for graphic visualisation of specific gene expression.

### Alternative splicing analysis

rMATS v4.1.2 (Shen et al., 2012; Park et al., 2013; Shen et al., 2014) was used to assess the alternative splicing events in each treatment, and to perform pairwise comparisons of alternative splicing events from the transcript count matrix. Like the DEG analysis, we compared the splicing events between M and N, N and Wt, Wt and NB1, NB2, and NB1 and NB2. Five types of events were considered: skipped exon (SE), alternative 5’ splice site (A5SS), alternative 3’ splice site (A3SS), mutually exclusive exon (MXE), and retained intron (RI). Using R (R Development Core Team, 2020), the package reshape (Wickham, 2007), and adapted methods from Tian and Monteiro (Tian and Monteiro, 2022), we selected the differentially spliced genes (DSG) with at least one differentially spliced site between treatments, a false discovery rate (adjusted p value) below 0.05 and a total junction count (for inclusion and skipping junctions) above 20. If a DSG had multiple spliced sites, we used the event with the maximum absolute value of inclusion level difference between the treatments (percent spliced in index, ΔPSI) to represent the differential splicing level of the gene. To select relevant DSG, we excluded low abundance splicing events likely to be splicing mistakes (|PSI|<50) (Tian and Monteiro, 2022). Plots including DSG were built using R (R Development Core Team, 2020), packages ggplot2 (Wickham, 2016) and wesanderson (Ram and Wickham, 2023). To visualize the gene junctions, we indexed the .bam files using samtools (Danecek et al, 2021), aligned the sequences with the genome (V2), and built the sashimi plots in the Integrative Genomics Viewer (IGV v2.17.3, Robinson et al. (2011)).

### Functional annotation

Gene descriptions were retrieved from the *Bicyclus anynana* genome annotation (Murugesan et al., 2022), and completed by locally blasting the assembly genes against the insecta (taxid:50557) NCBI non redundant (nr) protein database (built using the BLAST+ command line tool suite (Camacho et al., 2009)) in diamond v.2.0.8.146 (Buchfink et al., 2015; Buchfink et al., 2021). The alignment was imported in Blast2go v6.0.3 to get the GO annotations (Ashburner et al., 2000). Protein function prediction was done in parallel using InterProScan in Blast2go (Jones et al., 2014) and the eggNOG-mapper online tool (eggnog-mapper.embl.de (Huerta-Cepas et al., 2019; Cantalapiedra et al., 2021)). GO enrichment analyses were done with the R package clusterprofiler (Yu et al., 2012).

## Acknowledgment

We thank Jocelyn Wee, Suriya Murugesan and Shen Tian for their help with the RNAseq data analysis. We also thank Greenology for providing the corn plants. We acknowledge support from the Ministry of Education, Singapore (award MOE2018-T2-1-092), and the National Research Foundation, Singapore, Competitive Research Program (award NRF-CRP25-2020-0001).

## Funding

Ministry of Education, Singapore (award MOE2018-T2-1-092)

National Research Foundation, Singapore, Competitive Research Program (award NRF-CRP25-2020-0001)

## Authors contribution

Conceptualization: ED, AM

Investigation: ED, YPT, DZ

Funding acquisition: AM, ED

Project administration: AM

Supervision: ED, AM

Writing – original draft: ED, YPT

Writing – review & editing: ED, YPT, AM

## Notes

Authors declare no conflict of interest.

### Competing Interest Statement

The authors have declared no competing interest.

