## Supplementary material for "Butterfly brains change in morphology and in gene splicing patterns after brief pheromone exposure"

**Supplementary Materials, Dion et al.**

Supplementary Figure 1: Colour coded reference key for glomeruli identification
Supplementary Figure 2: RNAseq data summary
Supplementary Results 1: Description of relative positions and identification of glomeruli in the antennal lobe

**Supplementary Files:**
Supplementary File 1. Identification and volumes of glomeruli
- Table 1: Reference key for identifying individual glomerulus in the AL.
- Table 2: Total antennal lobe and glomeruli volumes of naïve females and males
- Table 3: Presence (Y) of individual glomerulus in each individual across all treatments used for analysis
- Table 4: Average relative volume (%) of the ten largest glomeruli for naïve females and males
- Table 5: Sizes and statistical analyses comparing all identified glomerulus of each individual across all treatment/group comparisons

Supplementary File 2: RNAseq summary and data description
- Table 1. Read depth and alignment rates
- Table 2. Number of DEG and DSG in each comparison
- Table 3. Output from the differential splicing analyses
- Table 4. Gene-set-enrichment analysis results for the DSG
- Table 5. Olfaction-related sensory genes present in the brain transcriptome
- Table 6. DSG with common significant |PSI| and splicing events between the treatment comparisons (associated with exposure to MSP2 changes)

File S3: List of DEG and DSG in all comparisons
- Table 1. Description and legend
- Table 2. MvsN
- Table 3. NvsWt
- Table 4. WtvsNB1
- Table 5. WtvsNB2
- Table 6. NB1vsNB2

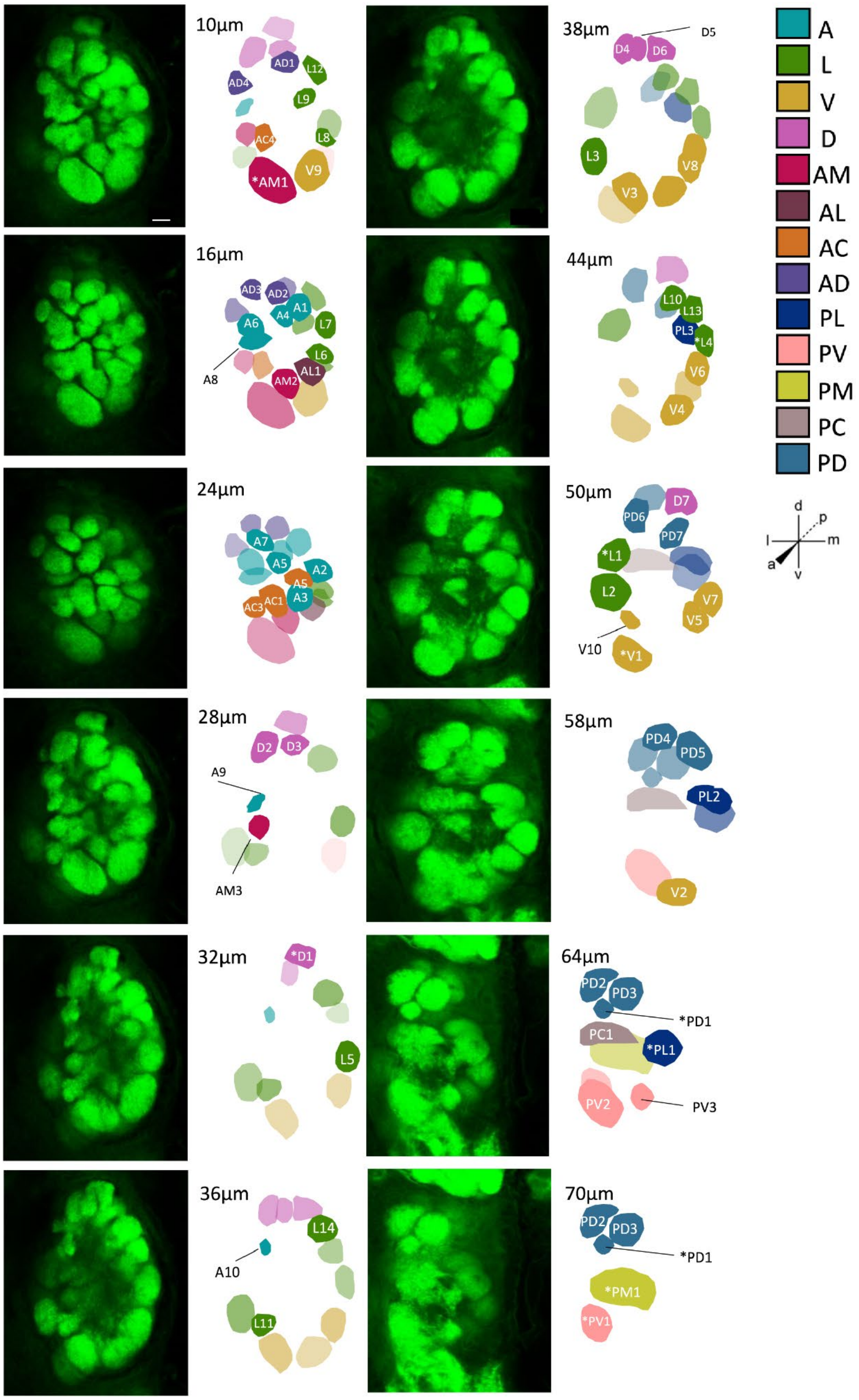


**<u>Supplementary Figure 1.</u> Colour coded reference key for glomeruli identification.** Glomeruli are coded and coloured based on their spatial group and subsequently numbered. Slices are imaged from the anterior to the posterior of the brain, with the depth of each slice indicated next to the image. Scale bar (white, top left image), 15 µm. * indicates a landmark glomerulus. Detailed description of glomerulus position is in Supplementary Results 1. A= anterior, L=lateral, V=ventral, D=dorsal, AM=antero-medial, AL=antero-lateral, AC= antero-central, AD=antero-dorsal, PL=postero-lateral, PV=postero-ventral, PM=postero-medial, PC=postero-central, PD=postero-dorsal. Orientations with axis indicate a, p, d. v. l .m for anterior, posterior, dorsal, ventral, lateral, medial. See Supplementary Results 1 below for details on identification and position of glomeruli.

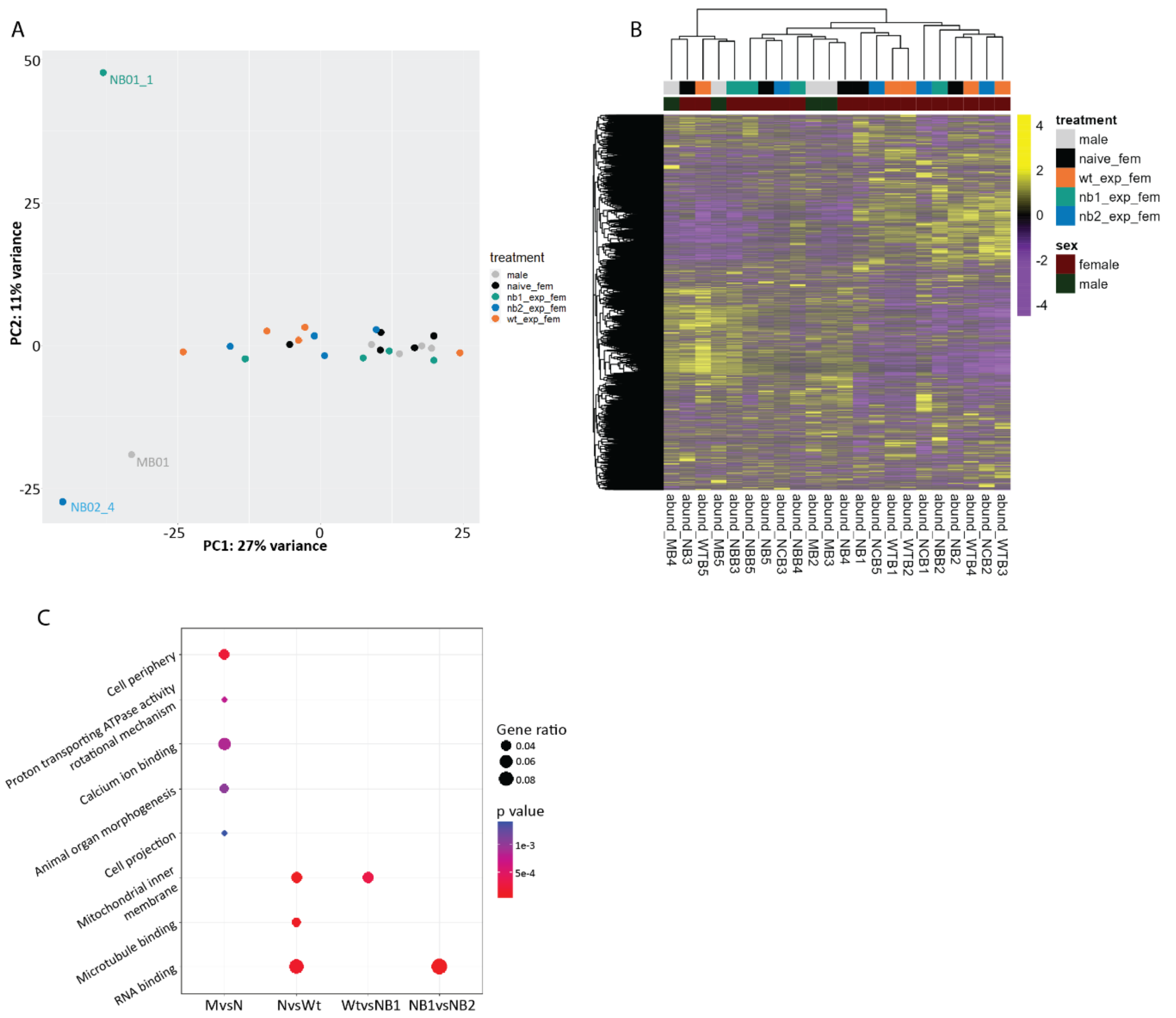


**Supplementary Figure 2. RNAseq data summary: There are more splicing events than significant changes in gene expression between treatments.** A) PCA plot representing expression data clustering built on 500 randomly picked genes from the count matrix; B) Hierarchical clustering heat map based on gene expression levels C) Gene-set-enrichment plot for the DSG showing the enriched key functions of differentially spliced genes in the comparisons.

**Supplementary Results 1. Description of relative positions and identification of glomeruli in the antennal lobe. Descriptions below apply to the right AL when imaged as in supplementary Figure 1.**

*Glomeruli in the Ventral Group.* There are 11 glomeruli partially located in the ventral region of the AL. Glomeruli V1 and V2 serve as landmarks. V1 is a relatively large glomerulus located at the very ventral left of the AL and appears towards the posterior end of the AL. V3 is relatively large, located antero-dorsally to V1 and V10 which is relatively small is located dorsal to V3. V2 is also a relatively large, smaller than V1 glomerulus located at the ventral right of the AL and appears towards the posterior end of the AL. V4 is a relatively large glomerulus and is located antero-dorsally to V2. V5, 6 and 7 are all similarly sized to V4. V5 is located postero-dorsally to V4.V6 is dorsal to V5, and V7 is posterior to V6. V8 is medium-sized and is located towards the anterior of V6, while V9 is a relatively large sized glomerulus and is located antero-medial of V8.

*Glomeruli in the Dorsal Group.* Nine glomeruli are located in the dorsal region of the AL. Glomerulus D1 serves as the landmark glomerulus and appears near the mid depth of the AL. D2 is a relatively large glomerulus that is located laterally to D1. D4 is a medium-sized glomerulus located towards the postero-dorsal side of D2. D5 is a medium to small sized glomerulus located medial to D4. D3 is a small glomerulus located antero-ventrally to D1 while D6 is a medium sized glomerulus located postero-dorsally to D1 and D7 is a relatively large glomerulus located medially to D6.

*Glomeruli in the Lateral Group.* There are 18 glomeruli located spatially in the lateral region of the AL with L1 and L4 being the landmark glomerulus. L1 is relatively large, near the posterior left end of the AL. L2 is located ventral to L1 and is a relatively large sized glomerulus. L3 is a medium sized glomerulus and is antero-ventral to L2. L11 is a medium to small sized glomerulus located on the medial of L3. L4 is a medium to small sized glomerulus, appearing near the posterior right end of the AL. L5 is a medium sized glomerulus and is located posterior to L4. L6 is located antero-dorsal to L4 and is a small sized glomerulus. L7 is a medium to small sized glomerulus located dorsally to L6 and L9 is a small glomerulus located medial to L7. L8 is a small sized glomerulus located antero-ventral to L5. Glomerulus L10 is located ventral to L4 and is a medium sized glomerulus. L12, L13 and L14 are all medium sized glomerulus, with L12 located to the dorsal side of L7, L13 to the postero-ventral side of L12, while L14 is located on the postero-dorsal side of L12.

*Glomeruli in the Anterior Group.* As there were no glomeruli in this region that were easy enough to distinguish at first glance, we decided to use a previously identified glomerulus, L9 as a landmark to identify the rest of the anterior glomeruli. There are 10 glomeruli in the anterior region of the AL. A1 is a medium sized glomerulus located antero-medially to L9. A2 and A3 are both medium-sized glomeruli located ventrally to A1 and A2 respectively. A4 is a small to medium sized glomerulus located medially to A1 and A5 is a medium sized glomerulus located dorsally to A4. A6 is a small to medium sized glomeruli located laterally to A4 while A7 is a medium sized glomerulus located dorsally to A6. A8 is a medium sized glomerulus that is located posterior of A6. A9 and A10 are small glomeruli with A9 located posteriorly of A8 and A10 located postero-dorsal of A9.

*Glomeruli in the Antero-Medial Group.* There are three glomeruli located in the anterior medial region of the AL. AM1, a relatively large glomerulus, serves as the landmark for this region and appears near the anterior of the AL. AM2 is a medium sized glomerulus located medial to AM1, and AM3 is a small to medium sized glomerulus located anterior-dorsally to AM1.

*Glomeruli in the Antero-Lateral Group.* There is only 1 glomerulus, AL1, which is located in the antero-lateral region of the AL. It was identified based on a previously identified glomerulus in the antero-medial region, AM2. AL1 is a medium sized glomerulus located laterally to AM2.

*Glomeruli in the Antero-Central Group.* There are four glomeruli located in the antero-central region of the AL. A previously identified glomerulus, A3, serves as the landmark glomerulus for this region. AC1 is a medium to small sized glomerulus located antero-medial to A3. AC2, 3, and 4 are all small sized glomerulus. AC2 is located dorsally to AC1, while AC3 is located lateral to AC1. AC4 is located lateral to AC1, and at the same time, dorsal to AC3.

*Glomeruli in the Antero-Dorsal Group.* There are four glomeruli located in the antero-dorsal region of the AL. Three previously identified glomeruli, A4, A5, and D2 serve as the landmark glomeruli for this region. All four glomeruli are small sized glomeruli, with AD1 located posterior to A4, AD2 postero-dorsal to A5, AD3 lateral to D2, and AD4 postero-lateral to AD3.

*Glomeruli in the Postero-Lateral Group.* There are three glomeruli in the postero-lateral region of the AL. PL1, a relatively large glomerulus, appears near the posterior end of the AL and serves as the landmark for this region. PL2 is also a relatively large glomerulus and is located antero-dorsally to PL1. PL3 is a medium sized glomerulus and is located anteriorly to PL2.

*Glomeruli in the Postero-Ventral Group.* There are three glomeruli in the postero-ventral region of the AL. PV1, a medium sized glomerulus appears near the posterior end of the AL, serves as the landmark glomerulus for this region, together with a previously identified glomerulus, V2. PV2 is a large sized glomerulus located ventrally to PV1, and PV3 is a small sized glomerulus located lateral to PV2.

*Glomeruli in the Postero-Medial Group.* There is only one glomerulus in the postero-medial region of the AL, PM1. This is a relatively large sized glomerulus that appears near the posterior end of the AL. As it is easily recognizable, it also serves as the landmark for this region.

*Glomeruli in the Postero-Central Group.* There is only one glomerulus in the postero-central region of the AL, PC1. It is identified through PM1, a previously identified glomerulus. PC1 is a medium sized glomerulus located dorsal to PM1.

*Glomeruli in the Postero-Dorsal Group.* There are 8 glomeruli in the postero-dorsal region of the AL, however one of the glomeruli, PD8, rarely appears throughout the sampled individuals. PD1 is a small sized glomerulus appearing near the posterior end of the AL and serves as the landmark glomerulus for this region. PD2 is a large glomerulus lateral to PD1, and PD4 is a medium sized glomerulus dorsal to PD2. PD3 is a relatively large glomerulus lateral to PD1, PD5 is a medium to large sized glomerulus dorsal to PD3, and PD7 is a medium sized glomerulus ventral to PD5. PD6 is a relatively large glomerulus anterior to PD2, and finally PD8 is a medium to small sized glomerulus anterior to PD4.
